# Operate a Cell-Free Biofoundry using Large Language Models

**DOI:** 10.1101/2024.10.28.619828

**Authors:** J. Hérisson, A. N. Hoang, A. El-Sawah, M. M. Khalil, J.-L. Faulon

## Abstract

In this paper, we present a novel approach to optimizing cell-free protein synthesis (CFPS) systems using artificial intelligence (AI), specifically leveraging ChatGPT-4 for code generation and active learning (AL). This study aims to automate and enhance the process of producing antimicrobial proteins, namely colicin M and colicin E1, in CFPS systems. We developed an automated workflow that employs an iterative Design-Build-Test-Learn (DBTL) cycle, integrating a newly implemented AL method with cluster margin (CM) selection to efficiently explore experimental conditions. The workflow components, including modules for sampling, plate design, instruction generation, and data analysis, were coded using ChatGPT-4 without further human modification. By employing this automated approach, significant improvements in protein yields were achieved, with a 9-fold increase for colicin M and a 3-fold increase for colicin E1 compared to standard buffer compositions. The use of LLMs in conjunction with AL demonstrated the potential of AI-driven methodologies to accelerate the optimization of complex biological processes and reduce manual intervention. The study also discusses limitations such as variability in CFPS and suggests future improvements in automation, reproducibility, and integration of diverse liquid handling systems to further enhance the scalability and efficiency of cell-free biofoundries.

## INTRODUCTION

Proteins are naturally produced in all cells to carry out a wide range of biological functions. Controlling their production can meet needs across various societal sectors, including health, environment, energy, and food. Since the advent of synthetic biology about twenty years ago, it has become relatively easy to modify a living cell to produce a desired protein. However, cellular production comes with significant challenges, especially regarding the integrity of the cell. For the past decade, it has been possible to overcome these cellular constraints by using minimal cell-free systems, which contain only the essential components needed for protein production. Industrializing these cell-free systems would greatly help in addressing the challenges of today and tomorrow, and cell-free protein synthesis systems (CFPSs) are now well established into robust platforms not only to synthetize a wide variety of compounds but also bioactive ones^1–4^. The increasing availability of commercialized cell-free and substrates renders it easier to make a CFPS platform accessible to any research laboratory. However, its widespread adoption is hampered by the need to further develop various tools to streamline and automatize the process. This encompasses the ability to accurately assess protein yields, select an appropriate acellular matrix, and optimize assembly of CFPS components^1,4,5^.

Over the last decade, in addition to advances in cell-free techniques, synthetic biology has seen its methods becoming automated within platforms. These platforms, not limited to any specific equipment, bring together in a unique place all the necessary tools to carry out a project. In 2019, a world-wide coordination has created the Global Biofoundries Alliance that currently encompasses more than 30 academic biofoundries all over the world^15^. Biofoundries are dedicated to advancing and expediting research in the fields of engineering and synthetic biology for both academic and practical applications. They support this by leveraging automation, high-throughput technologies, process upscaling, computer-aided design tools, and innovative methodologies. The use of iterative Design-Build-Test-Learn (DBTL) cycles in biological engineering empowers researchers to evaluate extensive genetic constructs and utilize artificial intelligence (AI) and machine learning (ML) to refine the design processes. Additional objectives involve fostering the growth of a sturdy engineering and synthetic biology sector and hastening the journey from conceptualization to market for biotechnological innovations and biomanufacturing engineering. A visionary aim is to devise biodesign principles that enable the effective modification of living cells for uses in biotechnology and medicine, offering a profound understanding of the intricacies of life. Nowadays, most of biofoundries have intensively developed the Build and Test steps with a large set of equipment and at a very good level of automation. Many manufacturers propose integrated solutions that can handle devices relative to the Build step (e.g. liquid handlers) as well as ones relative to the Test step (e.g. plate readers) through programmable features (workflows and scheduling) and robotic arms. However, if all biofoundries use pieces of software in their Design part, these are poorly automated, forcing users to convert data from one software tool to make them compatible with another. In addition, few of them use machine learning techniques, whether within tools or through the Learn step. Moreover, pieces of software developed in biofoundries are very sophisticated and are the result of several months of work by experts in both programming and the scientific field of the biofoundries.

To set up an efficient cell-free system, one roadblock is to find the optimal mix of different components from cell extract. This process needs to have a deep knowledge of both cell-free and components involved and can be time consuming and expensive. Moreover, as cell-free is a recent technology, there is not much data publicly available, noticeable fluctuation in components activities, therefore optimized production yield needs to be carried on for each specific system and for each protein of interest produced using these systems. To address this roadblock, different active learning (AL) frameworks have been adapted to search for a “global optimal” mix in a rather limited number of experiment trials^6,7^. AL is a machine learning approach where the model selectively queries the most informative data points for labeling to improve its performance efficiently. With an AL model, which requires a model to select informative input data, one can choose which data should be added for training more efficiently, achieving greater performance with less data.^8^. One of the most important aspects of the AL process is how much new data to label before retraining the model^9^. Classical AL methods assume that only one data sample is taken at a time (sequential AL), while retraining the model every time is inefficient. Moreover, the time required for retraining a model may not be neglected, or the addition of a single sample in the training set may not make a meaningful change, especially for deep learning models^10^. Therefore, in situations where multiple experiments are conducted in parallel, batch AL has been proposed as a solution that allows several data points to be labeled before retraining the model^11^. This approach is particularly effective because it enables multiple samples from the pool of data to be selected and labeled at the same time. However, sampling only based on uncertainty often results in bias, where the currently selected sample is not representative of the distribution of unlabeled datasets. In batch AL, it is essential for the selection criteria to account for both the diversity of the samples and the amount of information each sample provides relative to the model^12^. Nevertheless, focusing solely on diversity in sampling can increase labeling costs, as it may result in selecting multiple samples with low information content^13^. A very simple and easy-to-implement strategy has been proposed that considers both diversity and uncertainty of new data, called Cluster Margin^14^.

Large Language Models (LLMs) have revolutionized natural language processing (NLP) and shown significant potential in automating code generation. These models leverage vast amounts of textual data to understand and generate human-like text, including programming code or machine-readable protocols^16^. There are many ways to generate code from NLP, among the most famous ones we can cite ChatGPT, GitHub Copilot, Visual Studio Code… However, in most cases the code must be corrected or tuned by the programmer to run as expected. Beyond the code itself, LLMs can be also used to understand already written code^17^ or generate high-quality test sets^18^. Recently, ChatGPT has even been used to design, plan, and perform complex chemical synthesis experiments^19^ like browse the web, access documentation, control robotic systems, and collaborate with other LLMs to carry out six key tasks: planning chemical syntheses, navigating hardware documentation, executing cloud lab commands, controlling liquid handlers, managing complex multi-module tasks, and solving optimization problems using experimental data.

In this article, we leverage the power of AI to develop an automated workflow able to optimize a cell-free protein synthesis (CFPS) system. The workflow has been used to optimize the production of two antimicrobial proteins (namely colicin-M and colicin-E1). The code was fully written by ChatGPT-4 deploying a new AL method applied to reach a higher protein productivity.

## RESULTS

In this paper, we have engineered an automated protocol to enhance protein production in a cell-free system. Within the span of a mere week, this streamlined process is able to dramatically increase the yield of a cell-free system. This notable advancement is propelled by a very recent method of AL employing cluster margin (CM) selection criteria. Notably, except for the Learner module, all the computational code underpinning this study was autonomously generated by ChatGPT-4, with no subsequent human alterations. Below, we describe how the workflow works and how we used it to increase the production of colicin M and colicin E1 proteins by a factor of 9 and 3 respectively.

### Coding the workflow

Previously, a Python codebase using traditional programming methods to optimize a cell-free system was developed by our research group engineers. This task required about 2 months of full-time effort. Subsequently, we leveraged ChatGPT-4 to optimize and rewrite some parts of the code within a significantly shorter timeframe of one week. The outcome was a more concise code that effectively optimized the use of existing libraries. However, due to the complexity of the modules, the generated code had to be manually fixed or fine-tuned to incorporate additional features or tailor the results to their specific requirements. In the current work, we adopted a novel approach by directing ChatGPT-4 to autonomously write the entire code so that it is runnable without any subsequent human modifications. We proceeded module by module, using one different chat per module, and followed a step-by-step method. For each piece of code within a module, we first asked ChatGPT-4 to write a Python script that accomplishes a specific task based on the attached input data files, keeping our requests high-level without delving into programming details. Then, after copy-pasting and running the code, we asked ChatGPT-4 to fix errors (if errors appeared) or to add new features. At the end of this writing process, we proceeded the same way to automate the workflow shown in Figure 1 through a Snakefile.

**Figure 1:**
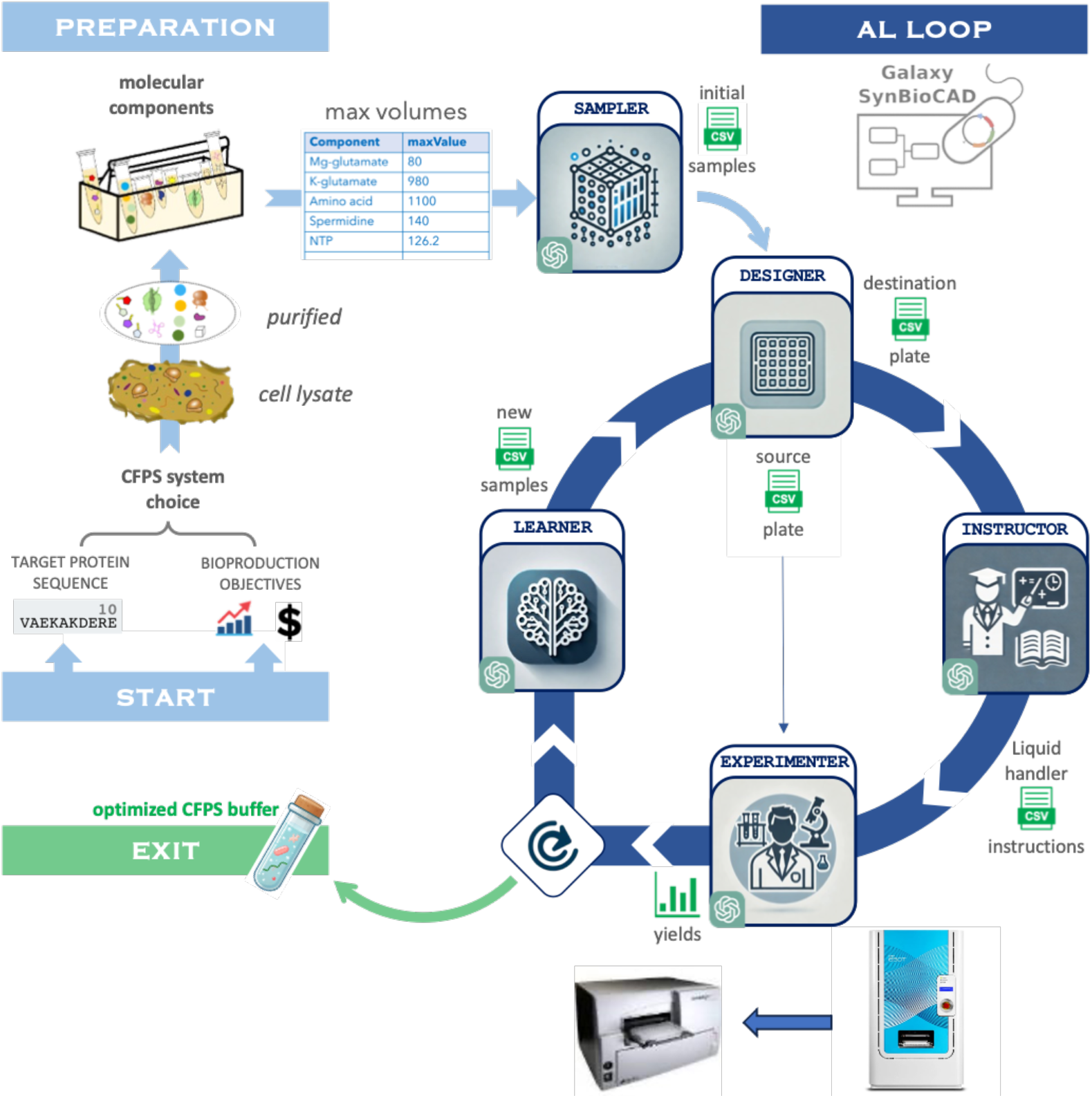
Active Learning loop. The AL loop searches for a mix with the highest yield by selecting and experimenting different cell-free recipes, then re-learning from its newly acquired data and suggesting the next experiments to test. The workflow starts by first using the SAMPLER once, then the loop begins with the plate DESIGNER used to prepare the source microplate and feeds the INSTRUCTOR which writes the instructions for an ECHO liquid handler. Both source plate and instruction file are used by the EXPERIMENTER that encompasses the liquid handler and a microplate reader to measure the fluorescence level of the destination plate. If the AL process is not finished, the fluorescence levels are analyzed and calibrated before feeding the LEARNER module which gives new samples to test. Once the model is well fitted, a last round is processed with the predicted best volume mixes to test.

The workflow is composed of different steps organized by an AL loop. It starts with an initialization step to provide the first cell-free compositions to test, then an active learning loop alternates between the software-based instructions and bench experiment. To initiate this workflow, the user inputs the maximum volumes of each cell-free component into the system. Thereafter, the AL loop commences, prompting the user to conduct a series of new experiments during each iteration. The user prepares the necessary materials and submits the results back to the Learning node. At the end of this process, which is marked by a substantial increase in production, the user is equipped with an optimized cell-free buffer tailored for the efficient synthesis of the desired protein.

#### The SAMPLER module

At the beginning of the workflow, the user inputs the maximum volumes of the components constituting the cell-free system and specifies the desired number of samples into the SAMPLER module. This module then generates an initial sampling of volume combinations to be tested. To ensure a representative subset of solution space, the module employs the Latin-Hypercube method for this initial sampling (refer to the METHODS section for details). The output is a CSV file containing these sample combinations^a^.

#### The plate DESIGNER module

To streamline the process, the plate DESIGNER module converts the cell-free samples into an intermediate representation of source and destination microplates. It analyzes the volumes specified in the SAMPLER module’s output file to determine the required volumes for the source plate. The result is two CSV files detailing the component volumes distributed across the source and destination plates^b^. The experimentalist uses the source plate file to prepare the source microplate, which is then placed into the liquid handler.

#### The INSTRUCTOR module

The INSTRUCTOR module translates the destination plate data into ECHO instructions. This module formats the data to comply with the liquid handler’s specifications, considering file formatting requirements and limitations related to high volumes or component viscosity. Additionally, water is added to achieve the final sample volume specified by the user. The output is a CSV file containing the instructions for the liquid handler^c^.

#### The EXPERIMENTER module

The prepared source plate, based on the plate DESIGNER’s output file, is inserted into the ECHO 650 Liquid Handler robot. The instruction file automatically generated by INSTRUCTOR module is loaded into the robot-operating software “Cherry Pick’’ on the computer, enabling the volume transfers from the source to the destination plate to create each sample as designed by the plate DESIGNER module. The destination microplate is then sent to a BioTek Synergy microplate reader, which records the fluorescence of each well over a 24-hour time course, the data is stored in a CSV file. The split GFP system was utilized to facilitate monitoring of the production kinetics of the antimicrobial peptides, colicins M and E1. Finally, a piece of code generated by ChatGPT-4 analyzes and calibrates data to feed the LEARNER module.

#### The LEARNER module

Finally, the LEARNER module uses the protein production of each well (sample) to learn their components’ interaction and suggests new experiments to run. These recommendations are provided in a new sampling file (CSV) for further testing.

Each module is available as open-source code^d^, a Conda package^e^, and a tool in the Galaxy Tool Shed^f^. Additionally, all these modules can be executed independently via command-line interface (CLI) on different platforms (Linux, macOS, Windows) or as an automated workflow through a Galaxy web portal. This flexibility ensures that users can integrate these tools seamlessly into their existing workflows, enhancing the reproducibility and efficiency of their experiments.

### Using the workflow

We used the workflow to optimize the production of colicins M and E1, two antimicrobial proteins. Developing novel antimicrobial therapies has recently increased for fighting antibiotic resistance. To address this need, we used a CFPS prokaryotic system based on *E. coli* lysate to rapidly and efficiently produce colicins M and E1 by circumventing the complexities and limitations associated with the living cells. This approach facilitated the high-yield production of both colicins, in addition to the precise manipulation of reaction parameters to optimize their synthesis.

#### Preparation of the experiment

The experimental design was based on the composition of the reaction mixture. Our cell-free system comprised essential components including Mg- and K-glutamate, amino acids, spermidine, 3-phosphoglyceric acid (3-PGA) or creatine phosphate as the energy source, nucleoside triphosphates (NTPs), PEG-20, HEPES, two plasmid DNA constructs (GFP 1-10 and GFP 11 fused to colicin M or colicin E1) for co-expression, and a cell extract. Certain components, such as tRNA, coenzyme A (CoA), NAD, cAMP, and folinic acid, which have been shown to have a minimal impact on protein production, were excluded from the reaction mixture^6^.

To assess the productivity of the cell-free system for colicin M, we tested six concentration levels (0%, 20%, 40%, 60%, 80% and 100%), where 80% corresponds to the standard buffer composition^20^. All components were varied as parameters except for HEPES which was used to maintain the pH of the system, and the bacterial cell-free lysate. This resulted in a combinatorial space of 6^9^ possible cell-free buffer compositions. In the case of colicin E1, the parameters were varied based on a fixed volume variation range of 20 nL from the max volume that corresponds to the 125% of the standard buffer composition. This made the combinatorial space as 4 * 49 * 55 * 7 * 47 * 7 * 43 * 62 * 66 > 10^12^ possible compositions.

Creatine phosphate was used as the energy source for the GFP-colicin E1 system with a reference concentration^21^, and 3-PGA was used for the GFP-colicin M system. Mg- and K-glutamate were individually standardized through preliminary experiments testing various combinations specific to each cell-free system for colicin M and colicin E1 (details in the Methods and supplementary section).

#### All-in-one Sampler - Designer - Instructor modules

First, we converted the six concentration levels of each of the 9 components—excluding HEPES, which had a constant concentration—into corresponding volumes and organized them into a CSV input file. This file was then processed using the Galaxy-SynBioCAD workflow *AI-CellFree – Init*^g^, which integrates the SAMPLER, plate DESIGNER, and INSTRUCTOR modules, resulting in the generation of ECHO instructions.

The colicin M workflow was parameterized with the following settings:

- <u>SAMPLER</u>: Number of samples: 100; Ratios: 0, 0.2, 0.4, 0.6, 0.8, 1
- Plate <u>DESIGNER</u>: Plate dimensions: 16×24; Number of replicates: 4/5/6; Sample volume: 10.5 µL
- <u>INSTRUCTOR</u>: Maximum transfer volume: 1000 µL; Split threshold: 1000 µL; Split components: HEPES, K-glutamate, Amino acid

We thus obtained four ECHO instructions files: one for each split component mentioned above (to be transferred first for viscosity reason) and one for the rest of the components. In each file, volumes to be transferred from which source well to which destination well were specified.

Concerning colicin E1, we used another strategy where no components were split, however, efficient transfer was ensured by changing the ECHO transfer mode from GP3 to CP for the viscous and critical components. The workflow was set as follows:

- <u>SAMPLER</u>: Number of samples: 100; Step: each 20 nL from 0 to max volume
- Plate <u>DESIGNER</u>: Plate dimensions: 16×24; Number of replicates: 4/5/6; Sample volume: 10.5 µL
- <u>INSTRUCTOR</u>: Maximum transfer volume: 1000 µL; Split threshold: 1000 µL; Source plate type: 384PP_AQ_CP for K-glutamate, creatine phosphate, NTPs, GFP1-10, and GFP11_Col-E1

In this case, one ECHO instruction file was the output, which was used as the only input file for the ECHO CherryPick software to transfer the different volumes of the components from each source well to its specific well in the destination plate

Once the first loop was completed, after each Learn step, the workflow *AI-CellFree – Core*^h^ which encompasses plate DESIGNER and INSTRUCTOR was run.

#### The Experimenter module (product quantification)

The instructions file was loaded into the ECHO’s software and the specified volumes of the different components are transferred. Since transferring all the components with ECHO takes more than 3 hours, the remaining volumes of lysate and water were manually pipetted into each destination well using a multichannel pipette, reaching a final volume of 10.5 µl. Then the destination microplate was loaded into a microplate reader (BioTek) in order to measure the production level in each well of the proteins of interest. We utilized the split GFP system^22^ with GFP1-10 and GFP11 fragments to construct our new split GFP-colicins system (see Methods section).

A 7-fold increase was observed after the full-time course of 24 hrs of co-expression of the two constructs for the colicin M system (Figure 2a), and a 20-fold increase for the colicin E1 system compared to the GFP1-10 fragment expression (Figure 2b). The fluorescence readings of all samples were normalized to those of the no DNA control. The fluorescence of the reassembled constructs of the two plasmid DNA (GFP 1-10 and GFP 11 fused to colicin M or colicin E1) show a rapid increase in activity, reaching a plateau at approximately 10 hours, which indicates a higher efficiency in producing the fluorescent protein than that of the sfGFP positive control. The GFP1-10 construct shows a gradual increase in fluorescence, however, its level remains significantly lower compared to the reassembled GFP. The GFP11-colicin M and GFP 11-colicin E1 fragments and the No DNA sample show relatively stable, minimal, almost no fluorescence production as expected.

**Figure 2:**
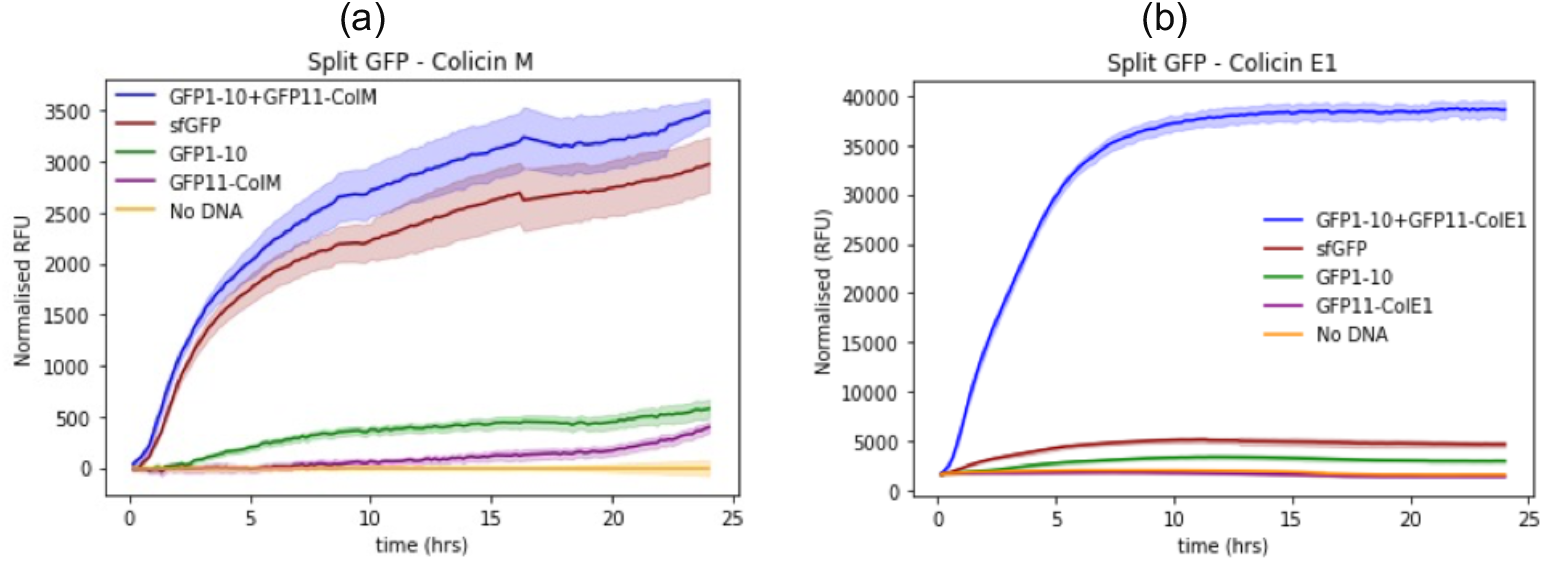
Kinetics of fluorescence measurements for the two split GFP-colicin systems. The graph shows the normalized relative fluorescence units (RFU) over 24 hours using DNA constructs with final conc. of 6 nM for GFP-ColM and 5 nM for GFP-ColE1. (a) In the split GFP-ColM system, the fully reassembled construct, shown in blue, has the highest fluorescence and is similar to the sfGFP control in red. (b) The split GFP-ColE1 system indicates a successful reassembly of the GFP with very high activity, shown in blue. In both figures, the other constructs act as controls to determine autofluorescence.

#### The Learner module

Because there is quite high variability in cell-free systems, a calibration process (see Methods) was needed to standardize the fluorescence levels of one plate in relation to the others. Then, the calibrated data were processed by the Learner module to train the model incrementally. The Cluster Margin (CM) was used to train the model better by using fewer data points (see Methods section). At each loop, the Learner module took the n-replicated (n = 3, 4 or 5) 50 sample levels to train the model and gave new samples to initiate the next loop toward the plate Designer module. Figure 3 shows that after four loops, we obtained a 9-fold yield increase (∼ 5-fold in average) for colicin M, while a 3-fold yield increase (∼ 2-fold in average) for colicin E1, compared to the yield obtained from the standard buffer found in literature^20^. During the first loops, our focus was not solely on finding the highest results but also on including some lower outcomes that provide valuable information about the relationship between components and yields. This approach helped the model learn more effectively, and our Expected Improvement (EI) measurement helped balance these two objectives. As we prioritized yield optimization, the average yield increased from one loop to another.

**Figure 3.**
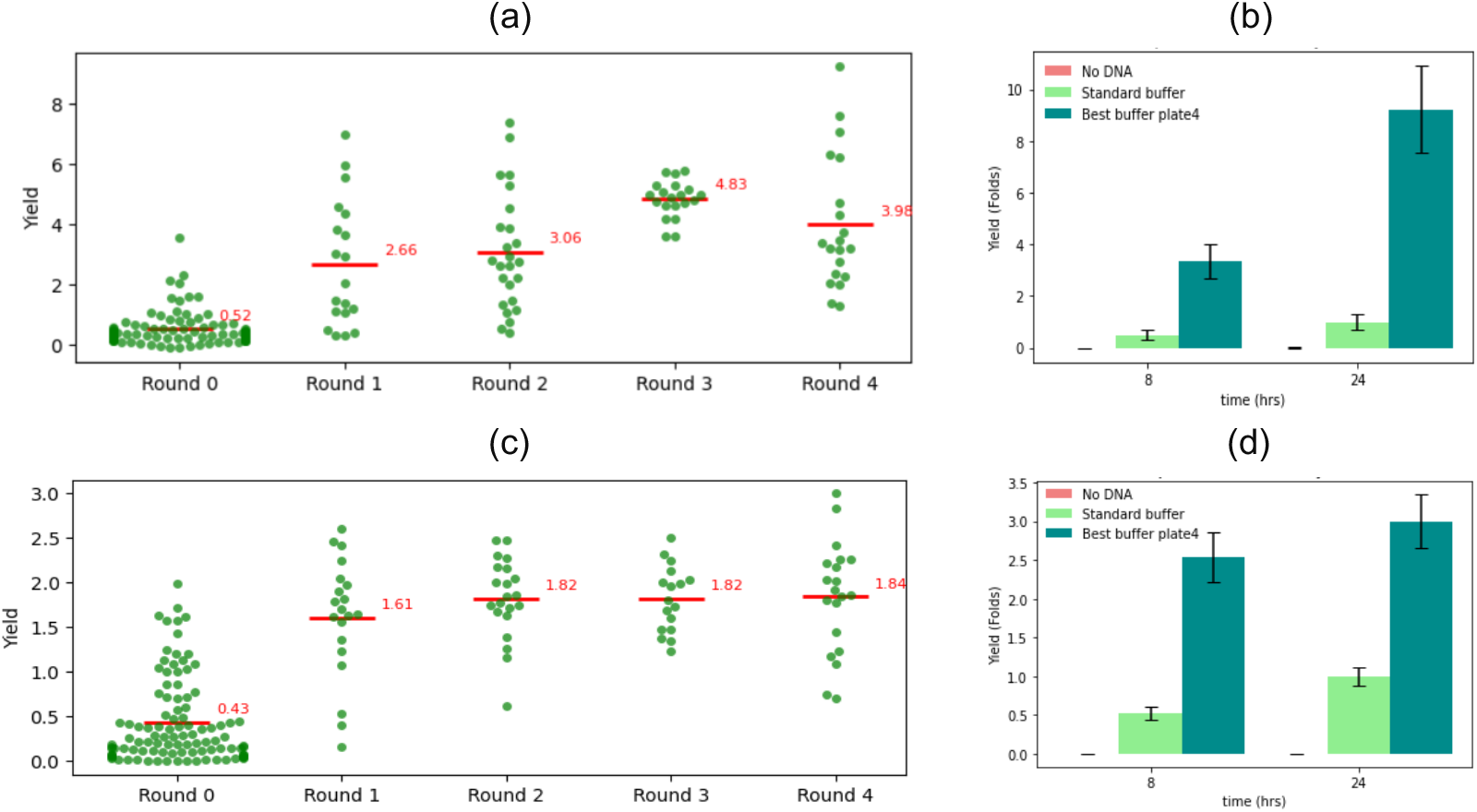
Comparison of achieved yields. The graphs (a) and (b) relate data for colicin M and the graphs (c) and (d) relate data for colicin E1. The swarm plots compare the yields achieved over four loops of testing using cluster margin active learning strategies. Each round (or loop) shows the average yields marked by the horizontal line in red. The data indicates the distribution and variability of yields for each round, illustrating the improvement of our bacterial cell-free system for both colicin M (a) and E1 (c). The graphs (b) and (d) show the maximum yield fold change of the best predicted buffer compared to the standard buffer composition across two time points (8 and 24 hours) that ensure a doubled yield for the standard buffer. The data show a very clear improvement between the last round (Plate 4) and the standard buffer, for both colicin M (9x) and E1 (3x).

In Loop 0, component combinations were generated using the Latin-Hypercube Sampling (LHS) method from the Sampler module. At this stage, low yields were observed (Figure 3a and 3c), likely due to the exploration-focused nature of the search. The median yields hovered around 1, however, some outliers suggested occasional higher-than-average yields, hinting at potential areas for improvement in subsequent loops. In Loop 1, a few higher yields were detected, although these were likely due to chance, given that early-stage models were still underperforming. Nevertheless, yields began to increase as the loops progressed, meaning the performance of the cell-free system improved, with average yields increased in Loops 2 and 3 (Figure 3a), and slightly increased in Loops 3 and 4 (Figure 3c). By Loop 4, although a few higher yields were achieved, it became evident that a yield plateau had likely been reached based on two key factors: first, the models began prioritizing exploration in regions associated with lower yields, which reduced the overall average; and second, model performance did not improve even as more data was added. The models became more confident in identifying high-performance regions, suggesting that the maximum possible yield within this search space had been reached. At the end, our model successfully predicted an optimized buffer that resulted in a 9-fold increase in yield for colicin M (Figure 3b) and a 3-fold increase for colicin E1 (Figure 3d), compared to the standard buffer composition.

## DISCUSSION

In this study, we successfully demonstrated the application of large language models (LLMs), specifically ChatGPT-4, to automate the optimization of protein production in a cell-free system using an active learning (AL) approach. By utilizing the ready-to-use Python code generated by ChatGPT-4, we were able to significantly reduce the time and complexity involved in programming tasks, from a couple of months to one week. Moreover, all pieces of code have been integrated into the workflow manager Galaxy. Thus, it is quite easy to install the developed tools on any instance of Galaxy. This innovative approach to leveraging LLMs for biological experimentation and to make tools available into Galaxy opens new avenues for integrating in the context of biofoundries.

One of the key achievements of our study is the development of an active learning loop, which iteratively improves cell-free system performance by identifying optimal combinations of components. The Cluster Margin (CM) selection criterion allowed for the exploration of diverse component mixtures while maintaining a balance between exploration and exploitation of high yield combinations and reducing the number of required experimental trials. At most 5 CM rounds were sufficient to multiply by 9 the colicin M yield compared to baseline and by 3 the one of colicin E1. This outcome illustrates the potential of AI-driven methodologies in optimizing complex biological systems where traditional methods are often time-consuming and data-intensive. Our iterative LLM driven DBTL cycle, combined with the AL approach could be broadly applied in synthetic biology to optimize various biological systems beyond cell-free protein synthesis.

Despite the success of this workflow, there are limitations and areas for improvement. The variability inherent in cell-free systems, such as fluctuations in component activity and lysate instability, necessitates careful calibration and standardization across experiments. Furthermore, part of this variability may stem from the inherent complexity of using two separate DNA constructs that must assemble to form the functional protein. Variations in folding or interaction dynamics between the expressed proteins can affect how well the constructs align and assemble, leading to differences in reassembly across replicates. Future work could focus on further automating the wet-lab steps, including sample preparation and data collection, to minimize human intervention and reduce experimental variability. Additionally, while our model converged on a very good solution within a few iterations, there may be potential to improve accuracy and yield further with additional iterations or by refining the sampling strategy used in the initial phase.

Looking forward, several areas of improvement can further streamline and enhance the automation of cell-free systems. One promising avenue for future development is addressing the reproducibility of CFPS systems. Unlike other biological systems, CFPS often exhibits higher variability due to fluctuations in component activity, batch-to-batch differences in lysates, and environmental factors. This variability can make it difficult to replicate experiments across different laboratories or even within the same lab over time. Based on numerical methods, future work could focus on identifying and mitigating the sources of this variability by identifying parameters of experiment that impact the result, which is critical for establishing standardized protocols in CFPS. Increasing the reproducibility of CFPS systems can help reduce the experimental costs and time required for large-scale protein production, making it a more reliable tool in both research and industrial applications.

Another key potential development is the integration of additional types of liquid handlers beyond the ECHO system used in this study, and the incorporation of robotic arms to automate more of the experimental process. Handling a variety of liquid handlers, including more widespread and accessible models, would increase the applicability of the workflow across different laboratories and platforms. By supporting a wider range of liquid-handling devices, the system could adapt more flexibly to diverse experimental setups. By introducing robotic arms for tasks such as sample preparation, microplate manipulation, and loading/unloading of liquid handlers and plate readers, the workflow could achieve a higher degree of automation, minimizing human intervention and contributing to reducing experimental variability and manipulating errors, and accelerate the optimization process, making the system more scalable for high-throughput applications.

As AI continues to evolve, we anticipate that future developments will lead to even more efficient, scalable, and autonomous systems, driving advancements in biomanufacturing and other areas of biotechnology.

## METHODS

### Preparation

#### DNA Constructs

The sequences for colicins M and E1 (Supplemental Table 1) were synthesized by GeneCust and inserted into pUC57 vectors. These were subsequently cloned into pIVEX vectors. PCR amplification of the colicin sequences and pIVEX backbone was performed using the primers provided in the Supplemental Table 2 and then digested by BsmBI and BbsI for a golden gate assembly. Similarly, the GFP 1-10 and GFP 11 fragments were cloned into pIVEX vectors, following the same procedure as for the colicins. Then, GFP 11 was used as a tagging system for colicins M and E1 by using BbsI to digest the pIVEX vector containing GFP 11, and BsaI was used for the colicin sequences (see details in Supplemental Table 3). All restriction enzymes used were sourced from NEB.

After each digestion, a DpnI treatment (NEB #R0176L) was performed for 1 hour to degrade any residual template DNA, followed by purification using the NEB PCR purification kit (NEB #T1130). The purified fragments were then ligated using T4 DNA ligase (NEB #M0202) according to the manufacturer’s protocol. Following ligation, the recombinant plasmids were transformed into chemically competent *E. coli DH5α* cells.

#### Plasmids preparation

The 3 plasmids for the GFP 1-10, GFP11-colicin M, and GFP 11-colicin E1 for the 2 systems were prepared at enough for all rounds of experiments. The plasmids were isolated from a 300 mL LB of *E. coli DH5α* using the Plasmid DNA purification NucleoBond Xtra Maxi of Macherey-Nagel. The plasmid pellets were resuspended in 1 ml and stored at -20 C. They were pooled to reach a concentration of ∼50 nM to be able to be transferred by Echo and then aliquoted for all the round/loop experiments avoiding any variation due to different batches.

#### Cell-free mix preparation and reactions

The cell-free components were prepared in sufficient batches/quantities for all experimental iterations and at stock concentrations optimized to be transferred by Echo 650 liquid handling system, ensuring the maximum volumes from all components together did not exceed the total reaction volume (10.5 µL) and were not below 20 nL for each component separately. All the reagents were prepared in enough amounts for all the experiments for the 2 systems of colicin M and E1 except for the 3-PGA (Sigma Aldrich) was used for colicin M while creatine phosphate (Sigma Aldrich) was used in the case of colicin E1.

The bacterial cell-free lysate was provided by Synthelis Biotech company, and the reactions were performed in 10.5 μL volumes at 30 °C in a 384-well plate. All the cell-free reaction components were set to be the system parameters altering at different concentrations/ratios except the lysate at 33% of the reaction, and HEPES at 50 mM as buffering reagent.

The components that were determined to be transferred by Echo underwent a drop test to determine the correct calibration to ensure uniform droplets at the center of the destination wells. The 384PP (PP-0200) plate was chosen as the source plate, while the calibration transfer mode was 384PP_AQ_GP3 for all reagents except for K-glutamate, NTPs, and DNAs for their viscosity using 384PP_AQ_CP mode. Cell lysate was added to the 384-well destination plate (Nunc 384-well optical bottom plates, Thermo-Scientific #242764) using a multichannel pipette after the transfer of all the other components by Echo.

As this study focused on optimizing the production of colicin M and E1 by varying the concentrations of the TX-TL buffer components to achieve the highest possible yield. We used the concentrations specified by Sun et al. as our 100% concentration level for all TX-TL components, which served as the standard/reference point. For the two plasmid DNAs of the GFP 1-10 and the GFP11-colicin M system, we set the concentration to 6 nM, while for the GFP 1-10 and the GFP11-colicin E1, it was set to 5 nM. These values represent the maximum achievable concentration within our reaction volume trying to maximize production from the standard buffer composition, representing a yield of 1.

After generating the instruction file, the source plate is filled based on the source plate map with the necessary reagents. The Echo cherry-pick application is then used to select the appropriate source and destination plates, and to upload the instruction file to initiate the transfers. During this process, exception errors may occur and should be promptly resolved to ensure the successful transfer of all reagents in the specified volumes.

### The Sampler

To create the different mixes, we needed to transfer volumes of solution from a source well-plate to a destination well-plate. The source plate is filled by hand while the destination plate is filled with a liquid handler, namely an ECHO 650. Because this device can handle volumes multiple of 2.5nL, we consider only such values for volumes to test. Therefore, discrete ranges must be considered as solutions space for each component, making it a finite set of solutions while remaining very large (e.g. millions of possible solutions for a few components). We can now generate a set of values to test within the component’s discrete ranges. Since we must handle duplicates or triplicates, generating 100 samples to test at each AL loop seems to be a good size. As such a subset is very small compared to millions of possible values, random values are not representative of the solution space since two random values can be close to each other. Therefore, because the quality of the first tested values can have a dramatic impact on the efficiency of AL, we preferred instead to use Latin-Hypercube Sampling (LHS) that provides a representative sampling of the full solution space^23^.

Then, this module takes as input a file containing the list of components in the cell-free system with their possible maximum volume and provides a LHS. To write this code, we use ChatGPT-4 (Data Analyst customized GPT) by feeding it with the input file (cf. Supplementary Material), asking the following first request:

*Write full commented and generic Python code that takes that kind of input file (header is column names) and produces a Latin-Hypercube Sampling in CSV file (with component names as header) where each line is a sample. The LHS will process on discrete ranges built from 0 to the maximum value (given by “maxValue column” in the input file) of each component (given by “Component” column in the input file) by increments of 2*.*5. Input and output files as well as the number of wanted samples will be passed in CLI arguments*.

After few iterations^i^, the code generated by ChatGPT-4 was executed to get the sampling file.

### The Designer

The conversion into ECHO instructions can be done from other data than a sampling of volumes. One would like to generate instructions from source and destination plate designed by hand or provided from another process. Then, we decided to implement another module that transforms sampling of volumes into source and destination plate. In the same way, we used ChatGPT-4 to code this module by giving the sample volumes file (CSV, cf. Supplementary Material) and the following initial request:

*Write a full commented and generic Python code that takes the sampling of volumes in this file and provides source and destination well-plates*.

*The program will take as CLI arguments:*

- *the sampling file (required)*
- *the wanted sample volume in the destination plate (required)*
- *for each of source and destination plate: starting well (default: “A1”), the plate dimensions (default: “16×24” for a 384 well-plate), the well capacity (default: 60000)*
- *for source plate only, the dead volume (default: 15000)*
- *the number of wanted replicates (default: 1)*
- *the output folder (default: current directory) to write source and destination files*.

*The destination plate is obtained by taking the sampling file and adding a first “Well” column with well labels in a vertical way (A1, B1*, …, *A2*…*), and a last column “Water” which is the volume of water to add to reach the wanted sample volume. Once the destination plate is filled, continue if the number of replicates is > 1*.

*The source plate also has the added columns “Well” (first) and “Water” last and is obtained by taking the sum of volumes for each component in the destination plate. The volumes are filled into one different well for each component*.

After some iterations^j^, the code generated by ChatGPT-4 was executed to get the source and destination files.

### The Instructor

This module takes as inputs source and destination plates and writes ECHO instructions. Similarly, ChatGPT-4 has been used to write the code by passing the source and destination files (CSV, see Supplementary Material) and asking initially for:

*Write a full documented Python code to write ECHO (liquid handler) instructions to transfer volumes from the source plate (source_plate*.*csv) to the destination plate (destination_plate*.*csv)*.

*Explore input file to know which format data have, but write a generic code*.

*The header of the output file has to be: “Source Plate Name,Source Plate Type,Source Well,Destination Plate Name,Destination Well,Transfer Volume,Sample ID” which are:*

- *“Source Plate Name” will be “Source[1]”;*
- *“Source Plate Type”, the type passed in CLI arg;*
- *“Source Well” is the well label where is the current component;*
- *“Destination Plate Name” will be “Destination[1]”;*
- *“Destination Well” is the well label where to drop the current component;*
- *“Transfer Volume” is the volume to transfer;*
- *“Sample ID” is the name of the component*.

*The CLI arguments will be:*

- *source plate file*
- *destination plate file*
- *output file*
- *source plate type for each component (default: “384PP_AQ_GP3”)*

After few iterations^k^, the code generated by ChatGPT-4 was executed to get the instructions file.

### The Experimenter

#### Fluorescence quantification (Split GFP)

Following the transfer of all reagents and the addition of cell lysate, the plate was incubated in a Synergy microplate reader at 30°C for 24 hours. The fluorescence kinetics were measured where the excitation wavelength was fixed at 485 nM, the emission at 528 nM and the gain at 50, using a slow linear shaking mode. The fluorescence was measured from the bottom of the 384-well plates sealed with multiwell plate sealers.

To measure our AMP production, we used the split-GFP system with GFP1-10 and GFP11 fragments. These fragments reassemble into a functional fluorescent protein, enabling the monitoring of gene expression in the E. coli TXTL system. colicin M is an AMP where its coding DNA sequence was fused to the GFP11 fragment and co-expressed with the GFP1-10 plasmid to allow for fluorescence quantification. This approach facilitated simple, real-time measurements of peptide production yield in TXTL, highlighting the significant advantage of the split-GFP system for screening and optimization procedures^24^.

The data generated from these measurements is subsequently processed to calculate the yield for each buffer, enabling further analysis and calibration. All fluorescence values were normalized to the autofluorescence value, which was determined using the same buffer composition without DNA, with water instead.

To compare different buffer compositions, we calculated the yield of AMP production based on the fluorescence values as the following:

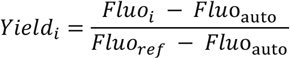

where:

- *Fluo*_*i*_ represents the measured fluorescence of the experimental composition,
- *Fluo*_*ref*_ denotes the fluorescence of the reference composition known to efficiently produce the AMP, and
- *Fluo*_*auto*_ accounts for the background fluorescence of the system without any GFP expression.

This equation allows for the normalization of fluorescence readings, enabling accurate comparison of colicins production yields across different compositions.

Through testing 329 buffer compositions for colicin M and 306 compositions for colicin E1, across four iterations excluding controls and replicates, we identified optimal conditions that significantly enhanced protein yield. Specifically, we achieved yields up to 9 times higher than the standard buffer for colicin M and 3 times higher for colicin E1, as illustrated in Figure 3.

### The Learner

#### Calibration

Due to cell-free variability, a calibration process was needed to standardize fluorescence levels of one plate in relation to the others. This step was completed with a piece of code written by ChatGPT-4 (CALIBRATOR). After iterating^l^, the code generated by ChatGPT-4^m^ was executed to calibrate date from one plate to another. The algorithm is described in the Supplementary Material.

#### Initialization

To define our search space, we took the maximum volumes^20^ for each component of the cell-free system as shown in Supplemental Table 4. Then, we initialized a design space based on the ranges of each component and selected an initial set of 100 experiments to run by using the Sampler module with the following command:

~~~
python -m icfree.sampler component_maxValues.tsv sampling.csv 100 --
seed 42 --fixed_values ‘{“HEPES”: 10}’ --ratios “0,0.2,0.4,0.6,0.8,1”
~~~

where:

- `component_maxValues.tsv` is the name of the file containing the maximum volumes of the cell-free components,
- `sampling.csv` is the name of the output file,
- `100` is the number of samples generated,
- `42` is the seed number for reproducibility purpose,
- `‘{“HEPES”: 10}’` is to fix the value of HEPES component to 10 μL,
- `“0,0.2,0.4,0.6,0.8,1”` is the list of the ratios to take as possible volume values for each of the components.

When the program ended, we obtained the 100 first cell-free combinations and we conducted this initial set of experiments to get the starting dataset.

##### Model Training

At initial state, a surrogate model was trained using Gaussian Process (GP) regression, using the first 100 measurements of fluorescence. GP is best-suited for optimization over continuous domains of less than 20 dimensions and our cell-free mix has 11 components^25^. In addition, GP can give uncertainty alongside each prediction, which is the core of informativeness estimation in AL. Then the model has been validated to ensure it accurately predicts system performance based on the components’ values.

##### Active Learning Loop

*Sampling*: Exploration the design space by iteratively selecting top few new points, Expected Improvement is used as an acquisition function to measure and rank the value of each new point to add into the model. This strategy is described in the next section.

*New Experimentation*: Conducting new experiments based on the selected points.

*Data Aggregation*: Adding the results of the new experiments to the existing dataset.

*Model Update*: Retraining of the surrogate model with the updated dataset.

*Convergence Check*: Assessing if the model performance or optimization goal meets a predefined convergence criterion (performance plateau if this work).

#### Sampling Acquisition Function

In this paper, we utilized Expected Improvement^26^ (EI) as an active learning acquisition function to select the top 50 most informative samples from a pool of 1000 randomly generated samples within a specified search space. This method is consistent with the Bayesian optimization framework for regression^25^, aiming to enhance model performance by identifying the best target and highest uncertainty, rather than using the margin as a metric for classification tasks^14^. The 1000 data points are ranked based on their EI value, with the highest values indicating the most informative samples for the model. After the 50 new points are labeled after lab experiments, they are added into the training set, and we retrain our GP regressor for the next loop.

#### Cluster Margin algorithm

The training process compared two methods: Vanilla Active Learning (VAL) and the Cluster Margin (CM) method. VAL selects the top 50 points from the search space based solely on their acquisition EI value, without considering the diversity of the data. In contrast, the CM method selects only 20 points, focusing on both informativeness and diversity. As training progressed, the R^2^ value increased significantly for both methods, indicating that each was successful in selecting informative data points for training (see Figure 4). Notably, while the CM method selected 60% fewer data points than VAL, it achieved similar model performance for the colicin M system, or even higher performance in the colicin E1 system. This demonstrates that increasing the amount of training data is not always necessary for model improvement. The traditional EI function is useful for evaluating the informativeness of individual data points but is less effective when applied to batch queries, where diversity plays a crucial role. The CM method overcomes this limitation by ensuring that selected data points are not overly like one another, resulting in a more diverse and representative dataset for training. This not only enhances model efficiency but also reduces resource consumption by minimizing the inclusion of low-performing samples. After four loops, the process was concluded, as further improvements in yield were not observed. However, it remains possible that additional iterations could increase the model’s accuracy.

**Figure 4:**
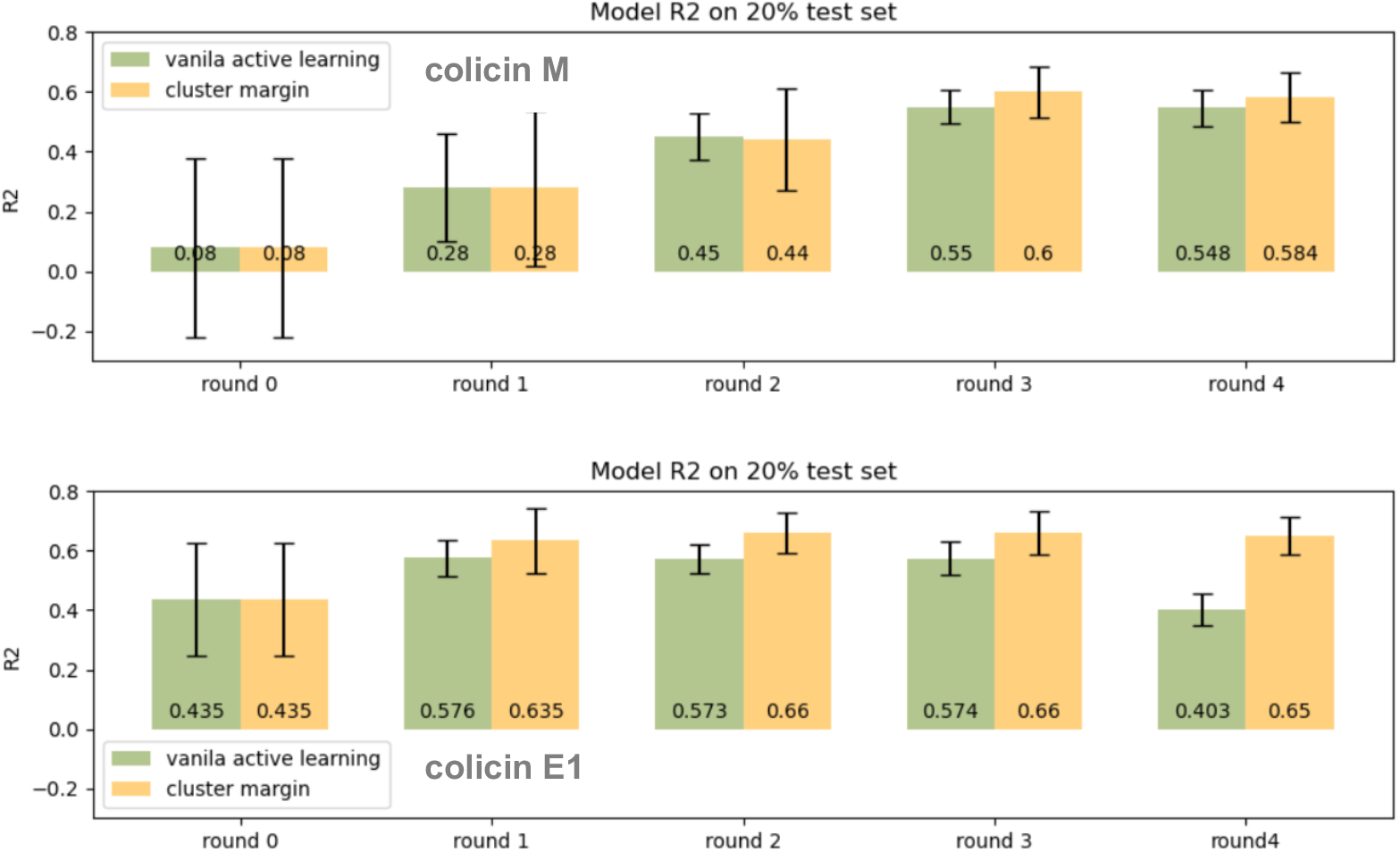
Comparison of model performances between vanilla active learning (VAL) and the cluster margin method (CM). Starting with the same initial dataset, different training sets are selected at each loop according to the respective methods. Consequently, two independent models are trained to suggest new data points. We observed the model R^2^ on a 20% test set and recorded the standard deviation over 100 repetitions.

To enhance more data representation through selected samples, we used the Cluster Margin method on top of the EI acquisition function. Initially, the entire dataset is segmented into k = 10 groups using Hierarchical Agglomerative Clustering (HAC), where samples within each group are considered sufficiently similar. This clustering method utilizes Average Linkage to measure distances between points, specifically avoiding the chaining effect associated with Single Linkage Clustering^27^. Single linkage clustering can force clusters together due to single elements being close, despite many other elements within each cluster being distant from each other. We selected a = 50 points using EI as a measurement like in Vanilla Active Learning (VAL), but following the selection of these examples, the clusters are utilized to pick the most diverse set of b = 20 points (where b < a) using a round-robin sampling strategy. These points are separated into their groups from HAC, and these clusters are then ordered by size, starting from the smallest to the largest. From the first cluster, the point with the highest EI is selected and repeated to the next bigger cluster until b points are chosen. This strategy ensures that the selected points are not only uncertain (as indicated by high EI) but also representative of different clusters or regions within the dataset.

## Supporting information

Calibration Algorithm

## ACKNOWLEDGEMENTS

Authors would like to acknowledge funding provided by the ANR funding agency grant numbers ANR-20-BiopNSE (ICFREE project), ANR-22-PEBB-0008 (PEPR B-BEST France 2030 program), ANR-21-ESRE-0021 (ALADIN project), and the UE HORIZON BIOS program (grant number 101070281). They also acknowledge M. Sabeti-Azad for her contribution in the plasmids design and her implication in this work. Finally, they thank Synthelis Biotech company, which is associated to the ICFREE project, for their contribution and the production of bacterial cell-free lysates.

## Footnotes

a https://github.com/brsynth/icfree-ml/blob/main/icfree/sampler.py

b https://github.com/brsynth/icfree-ml/blob/main/icfree/plate_designer.py

c https://github.com/brsynth/icfree-ml/blob/main/icfree/instructor.py

d https://github.com/brsynth/icfree-ml

e https://anaconda.org/bioconda/icfree-ml

f https://toolshed.g2.bx.psu.edu

g https://galaxy-synbiocad.org/u/joan/w/icfree-init

h https://galaxy-synbiocad.org/u/joan/w/icfree-core

i https://chat.openai.com/share/a6d8da15-3f78-434d-955e-efb7996ed07e

j https://chat.openai.com/share/56f141fe-0a97-412d-a782-39c2bf74aee6

k https://chat.openai.com/share/78f69117-75fb-4a51-83a7-813fd1ef0c98

l https://chatgpt.com/share/e/66e43b41-7330-8007-9d77-50a2e1f1c9ef

m https://chatgpt.com/share/e/67152c45-22b8-8007-9f72-cc160accea8e

## REFERENCES

1. Carlson, E. D., Gan, R., Hodgman, C. E. & Jewett, M. C. Cell-free protein synthesis: Applications come of age. Biotechnology Advances 30, 1185–1194 (2012).

2. Lu, Y. Cell-free synthetic biology: Engineering in an open world. Synthetic and Systems Biotechnology 2, 23–27 (2017).

3. Martin, R. W. et al. Development of a CHO-Based Cell-Free Platform for Synthesis of Active Monoclonal Antibodies. ACS Synth. Biol. 6, 1370–1379 (2017).

4. Pardee, K. et al. Portable, On-Demand Biomolecular Manufacturing. Cell 167, 248-259.e12 (2016).

5. Batista, A. C., Soudier, P., Kushwaha, M. & Faulon, J. Optimising protein synthesis in cell-free systems, a review. Engineering Biology 5, 10–19 (2021).

6. Borkowski, O. et al. Large scale active-learning-guided exploration for in vitro protein production optimization. Nat Commun 11, 1872 (2020).

7. Pandi, A. et al. A versatile active learning workflow for optimization of genetic and metabolic networks. Nat Commun 13, 3876 (2022).

8. Settles, B. Active Learning Literature Survey. (2009).

9. Rubens, N., Elahi, M., Sugiyama, M. & Kaplan, D. Active Learning in Recommender Systems. in Recommender Systems Handbook (eds. Ricci, F., Rokach, L. & Shapira, B.) 809–846 (Springer US, Boston, MA, 2015). doi:10.1007/978-1-4899-7637-6_24.

10. Kirsch, A., van Amersfoort, J. & Gal, Y. BatchBALD: Efficient and Diverse Batch Acquisition for Deep Bayesian Active Learning. Preprint at 10.48550/ARXIV.1906.08158 (2019).

11. Sugiyama, M. & Rubens, N. A batch ensemble approach to active learning with model selection. Neural Networks 21, 1278–1286 (2008).

12. Hino, H. Active Learning: Problem Settings and Recent Developments. Preprint at 10.48550/ARXIV.2012.04225 (2020).

13. Ren, P. et al. A Survey of Deep Active Learning. ACM Comput. Surv. 54, 1–40 (2022).

14. Citovsky, G. et al. Batch Active Learning at Scale. in Advances in Neural Information Processing Systems (eds. Ranzato, M., Beygelzimer, A., Dauphin, Y., Liang, P.S. & Vaughan, J.W.) vol. 34 11933–11944 (Curran Associates, Inc., 2021).

15. Hillson, N. et al. Building a global alliance of biofoundries. Nat Commun 10, 2040 (2019).

16. Jiang, S., Evans-Yamamoto, D., Bersenev, D., Palaniappan, S. K. & Yachie-Kinoshita, A. ProtoCode: Leveraging large language models (LLMs) for automated generation of machine-readable PCR protocols from scientific publications. SLAS Technology 29, 100134 (2024).

17. Nam, D., Macvean, A., Hellendoorn, V., Vasilescu, B. & Myers, B. Using an LLM to Help With Code Understanding. in Proceedings of the IEEE/ACM 46th International Conference on Software Engineering 1–13 (ACM, Lisbon Portugal, 2024). doi:10.1145/3597503.3639187.

18. Rao, N., Jain, K., Alon, U., Goues, C. L. & Hellendoorn, V. J. CAT-LM: Training Language Models on Aligned Code And Tests. Preprint at http://arxiv.org/abs/2310.01602 (2023).

19. Boiko, D. A., MacKnight, R., Kline, B. & Gomes, G. Autonomous chemical research with large language models. Nature 624, 570–578 (2023).

20. Sun, Z. Z. et al. Protocols for Implementing an Escherichia coli Based TX-TL Cell-Free Expression System for Synthetic Biology. JoVE 50762 (2013) doi:10.3791/50762.

21. Kim, T.-W.Kim, D.-M. & Choi, C.-Y. Rapid production of milligram quantities of proteins in a batch cell-free protein synthesis system. Journal of Biotechnology 124, 373–380 (2006).

22. Cabantous, S. & Waldo, G. S. In vivo and in vitro protein solubility assays using split GFP. Nat Methods 3, 845–854 (2006).

23. McKay, M. D., Beckman, R. J. & Conover, W. J. A Comparison of Three Methods for Selecting Values of Input Variables in the Analysis of Output from a Computer Code. Technometrics 21, 239 (1979).

24. Levrier, A. et al. Split minireporters facilitate monitoring of gene expression and peptide production in linear cell-free transcription-translation systems. Preprint at 10.1101/2024.05.16.594532 (2024).

25. Frazier, P. I. A Tutorial on Bayesian Optimization. Preprint at http://arxiv.org/abs/1807.02811 (2018).

26. Jones, D. R., Schonlau, M. & Welch, W. J. Efficient Global Optimization of Expensive Black-Box Functions. Journal of Global Optimization 13, 455–492 (1998).

27. Sun, Y. et al. A large-scale benchmark study of existing algorithms for taxonomy-independent microbial community analysis. Briefings in Bioinformatics 13, 107–121 (2012).

