## Supplementary material for "Operate a Cell-Free Biofoundry using Large Language Models": Calibration Algorithm

**Step 1: Yield Calculation**
Given fluorescence data matrix $F$ for the input file and reference file:

1. Let $f_i$ denote the fluorescence value for sample $i$.
2. Identify two rows for 'Jove+' and 'Jove-':
   - $J_+$: Average fluorescence value for the 'Jove+' row.
   - $J_-$: Average fluorescence value for the 'Jove-' row.
3. Calculate autofluorescence $A$ and reference value $R$ as:

$$A = \frac{1}{N}\sum_{j=1}^{N} f_{J-j}\,, \qquad R = \frac{1}{N}\sum_{j=1}^{N} f_{J+j}$$

   where $n$ is the number of fluorescence values.
4. For each sample $i$, compute the yield $Y_i$ as:

$$Y_i = \frac{f_i - A}{R - A}$$

**Step 2: Matching Components**

1. Given input file $I$ and reference file $R_f$, detect component columns $C$ before the fluorescence columns.
2. $\forall\, i \in I, \forall\, j \in R_f$, check if the rounded component combinations match:

$$round(C_i) = round(C_j)$$

**Step 3: Regression with Outlier Removal**

1. Let $Y_i$ and $Y_{ref,i} = aY_i + b + \varepsilon_i$ denote the average yield values for matching rows from $I$ and $R_f$, respectively.
2. Fit an ordinary least squares (OLS) regression:

$$Y_{ref,i} = aY_i + b + \varepsilon_i$$

   where $\varepsilon_i$ is the residual error.
3. Calculate Cook's distance for each point:

$$D_i = \frac{\hat{\varepsilon}_i^2}{p\hat{\sigma}^2}\left(\frac{h_i}{(1-h_i)^2}\right)$$

   Where $\hat{\varepsilon}_i$ is the residual for point $i$, $p$ is the number of parameters, $\hat{\sigma}^2$ is the variance, and $h_i$ is the leverage.
4. Iteratively remove the point with the highest Cook's distance until the $R^2$ value of the model exceeds a specified threshold $R^2_{limit}$, or up to 30% of the data points are removed.

**Step 4: Calibrated Yield**
For each computed yield $Y_i$, calculate the calibrated yield $Y_{calib,i}$:

$$Y_{calib,i} = aY_i + b$$

**Step 5: Picking of reference points for next batch** *(change reference points in order to cover all the range of yield values)*

- $R' \longleftarrow \emptyset$
- $R' \longleftarrow \{(x_{jove+},\ yield'_{jove+})\} \cup \{(x_{jove-},\ yield'_{jove-})\} \cup \{(x,\ yield'_{max})\}$
- $R' \longleftarrow R' \cup \{x \in_R S\}_{n-3}$
- Return $(S_{cal} \setminus R, R')$
