## Supplemental Table 1 for "Operate a Cell-Free Biofoundry using Large Language Models"

**Supplemental Table 1. Sequences for Colicins M and E1.** We used the sequences from *https://doi.org/10.1093/synbio/ysy004, found in the supplementary, but with excluding their modifications.*

**Colicin M sequence**

```
ATGGAAACCTTAACTGTTCATGCACCATCACCATCAACTAACTTACCAAGTTATGGCAATGGTGCATTTTCTC
TTTCAGCACCGCATGTTCCTGGTGCTGGACCTCTTTTAGTCCAGGTTGTTTATAGTTTTTTCCAGAGTCCAAA
CATGTGTCTTCAGGCTTTAACTCAACTTGAGGATTACATCAAAAAACATGGGGCTAGCAACCCTCTCACATTG
CAGATCATATCGACAAATATTGGTTACTTCTGTAACGCCGACCGAAATCTGGTTCTTCACCCTGGAATAAGCG
TTTATGACGCTTACCACTTCTCAAAACCAGCGCCAAGTCAATATGACTATCGCTCAATGAATATGAAACAAAT
GAGCGGTAATGTCACTACACCAATTGTGGCGCTTGCTCACTATTTATGGGGTAATGGCGCTGAAAGGAGCGTT
AATATCGCCAACATTGGTCTTAAAATTTCCCCTATGAAAATTAATCAGATAAAAGACATTATAAAATCTGGTG
TAGTAGGTACATTCCCTGTTTCTACAAAGTTCACACATGCCACTGGTGATTATAATGTTATTACCGGTGCATA
TCTTGGTAATATCACACTGAAAACAGAAGGTACTTTAACTATCTCTGCCAATGGCTCCTGGACTTACAATGGC
GTTGTTCGTTCATATGATGATAAATACGATTTTAACGCCAGCACTCACCGTGGCGTCATCGGAGAGTCGCTCA
CAAGGCTCGGGGCGATGTTTTCTGGTAAAGAGTACCAGATACTGCTTCCTGGTGAAATTCACATTAAAGAAAG
TGGTAAGCGATAA
```

**Colicin E1 sequence**

```
ATGGAAACCGCGGTAGCGTACTATAAAGATGGTGTTCCTTATGATGATAAGGGACAGGTAATTATTACTCTTT
TGAATGGTACTCCTGACGGGAGTGGCTCTGGCGGCGGAGGTGGAAAAGGAGGCAGTAAAAGTGAAAGTTCTGC
AGCTATTCATGCAACTGCTAAATGGTCTACTGCTCAATTAAAGAAAACACAGGCAGAGCAGGCTGCCCGGGCA
AAAGCTGCAGCGGAAGCACAGGCGAAAGCAAAGGCAAACAGGGATGCGCTGACTCAGCGCCTGAAGGACATCG
TGAATGAGGCTCTTCGTCACAATGCCTCACGTACGCCTTCAGCAACAGAGCTTGCTCATGCTAATAATGCAGC
TATGCAGGCGGAAGACGAGCGTTTGCGCCTTGCGAAAGCAGAAGAAAAAGCCCGTAAAGAAGCGGAAGCAGCA
GAAAAGGCTTTTCAGGAAGCAGAACAACGACGTAAAGAGATTGAACGGGAGAAGGCTGAAACAGAACGCCAGT
TGAAACTGGCTGAAGCTGAAGAGAAACGACTGGCTGCATTGAGTGAAGAAGCTAAAGCTGTTGAGATCGCCCA
AAAAAAACTTTCTGCTGCACAATCTGAAGTGGTGAAAATGGATGGAGAGATTAAGACTCTCAATTCTCGTTTA
AGCTCCAGTATCCATGCCCGTGATGCAGAAATGAAAACGCTCGCTGGAAAACGAAATGAACTGGCTCAGGCAT
CCGCTAAATATAAAGAACTGGATGAGCTGGTCAAAAAACTATCACCAAGAGCCAATGATCCGCTTCAGAACCG
TCCTTTTTTTGAAGCAACCAGACGACGGGTTGGGGCCGGTAAGATTAGAGAAGAAAAACAAAAACAGGTAACA
GCATCAGAAACACGTATTAACCGGATAAATGCTGATATAACTCAGATCCAGAAGGCTATTTCTCAGGTCAGTA
ATAATCGTAATGCCGGTATCGCTCGTGTTCATGAAGCTGAAGAAAATTTGAAAAAAGCACAGAATAATCTCCT
TAATTCACAGATTAAGGATGCTGTTGATGCAACAGTTAGCTTTTATCAAACGCTGACTGAAAAATATGGTGAA
AAATATTCGAAAATGGCACAGGAACTTGCTGATAAGTCTAAAGGTAAGAAAATCGGCAATGTGAATGAAGCTC
TCGCTGCTTTTGAAAAATACAAGGATGTTTTAAATAAGAAATTCAGCAAAGCCGATCGTGATGCTATTTTTAA
TGCGTTGGCATCGGTGAAGTATGATGACTGGGCTAAACATTTAGATCAGTTTGCCAAGTACTTGAAGATTACG
GGGCATGTTTCTTTTGGATATGATGTGGTATCTGACATCCTAAAAATTAAGGATACAGGTGACTGGAAGCCAC
TATTTCTTACATTAGAGAAGAAAGCTGCAGATGCAGGGGTGAGTTATGTTGTTGCTTTACTTTTTAGCTTGCT
TGCTGGAACTACATTAGGTATTTGGGGTATTGCTATTGTTACAGGAATACTATGCTCCTATATTGATAAGAAT
AAACTTAATACTATAAATGAGGTGTTAGGATTTAA
```
