## Supplemental Table 2 for "Operate a Cell-Free Biofoundry using Large Language Models"

**Supplemental Table 2. Primers used for PCR and the restriction enzymes.** *Each of these fragments of interest were first cloned into the pIVEX backbone plasmid. Colicins M and E1 were amplified from the pUC57 plasmid, using the same primers with the addition of the BsmB1 restriction sites, and then cloned into the pIVEX backbone using BbsI restriction enzymes. GFP1-10 and GFP11 fragments were amplified from plasmids already present in the lab.*

| Fragments | Primers used | For | Restriction Enzymes | Plasmid Backbone |
|---|---|---|---|---|
| Colicins (M or E1) | CcgtctcTTTAATACGACTCACTATAGGGAGACC<br>CcgtctcATCAGCAAAAAACCCCTCAAGACC | Fragment amplification | BsmBI<br>BsmBI | pIVEX |
| | GgaagacTTCTGAAAGGAGGAACTATATCCGG<br>AgaagacTATTAATTTCGCGGGATCGAGATC | Backbone amplification | BbsI<br>BbsI | |
| GFP1-10 | CcgtctcTATGTCTAAAGGTGAAGAACTGTTCACC<br>CcgtctcATCATTTTTCGTTCGGGTCTTTAG | Fragment amplification | BsmBI<br>BsmBI | pIVEX |
| | AgaagacAGACATGCGGCCTTCG<br>GgaagacTTATGATATCAAGATCCGGTAAGATCC | Backbone amplification | BbsI<br>BbsI | |
| GFP11 | CcgtctcTATGCGTGACCACATGG<br>CcgtctcATTATTTGTACAGTTCGTCCATACC | Fragment amplification | BsmBI<br>BsmBI | pIVEX |
| | AgaagacAGGCATGCGGCCTTCG<br>GgaagacTTATAATGATATCAAGATCCGGTAAGATCC | Backbone amplification | BbsI<br>BbsI | |
