## Supplemental Table 3 for "Operate a Cell-Free Biofoundry using Large Language Models"

**Supplemental Table 3. Primers used for adding GFP 11.**

| Fragments | Primers used | For | Restriction Enzymes | Plasmid Backbone |
|---|---|---|---|---|
| GFP11-Colicins (M or E1) | GgaagacTTCTAGCATAACCCCTTGGGG<br>AgaagacAGTTTGTACAGTTCGTCCATACCG | Fragment amplification | BbsI<br>BbsI | pIVEX |
| | AggtctcTCAAAAGGATCCGCATCGAAGG<br>AggtctcGctagTTATTGCTCAGCGG | Backbone amplification | BsaI<br>BsaI | |
