## Supplemental Table 4 for "Operate a Cell-Free Biofoundry using Large Language Models"

**Supplemental Table 4. Maximum volumes of the cell-free components.** *These values define the search space of our system, they represent the volumes in nL corresponding to the max. concentrations that we set to be 125 % of the reference concentrations.*

| Component | Max. Value |
|---|---|
| Mg-glutamate | 131.3 |
| K-glutamate | 350.0 |
| Spermidine | 131.3 |
| 3-PGA | 281.3 |
| NTP | 126.2 |
| HEPES | 262.5 |
| Amino acid | 1094 |
| DNA 1 | 1434.0 |
| DNA 2 | 1485.0 |
| PEG-8000 | 1312.0 |
